# Blood and Brain Gene Expression Trajectories Underlying Neuropathology and Cognitive Impairment in Neurodegeneration

**DOI:** 10.1101/548974

**Authors:** Yasser Iturria-Medina, Ahmed F. Khan, Quadri Adewale, Alzheimer’s Disease Neuroimaging Initiative

## Abstract

Neurodegenerative disorders take decades to develop and their early detection is challenged by confounding non-pathological aging processes. For all neurodegenerative conditions, we lack longitudinal gene expression (GE) data covering their large temporal evolution, which hinders the fully understanding of the underlying dynamic molecular mechanisms. Here, we aimed to overcome this limitation by introducing a novel GE contrastive trajectory inference (GE-cTI) method that reveals enriched temporal patterns in a diseased population. Evaluated on 1969 subjects in the spectrum of late-onset Alzheimer’s and Huntington’s diseases (from ROSMAP, HBTRC and ADNI studies), this unsupervised machine learning algorithm strongly predicts neuropathological severity (e.g. Braak, Amyloid and Vonsattel stages). Furthermore, when applied to *in-vivo* blood samples (ADNI), it predicts 97% of the variance in memory deterioration and its future declining rate, supporting the identification of a powerful and minimally invasive (blood-based) tool for early clinical screening and disease prevention. This technique also allows the discovery of genes and molecular pathways, in both peripheral and brain tissues, that are highly predictive of disease evolution. Eighty percent of the most predictive molecular pathways identified in the brain were also top predictors in the blood. The GE-cTI is a promising tool for revealing complex neuropathological mechanisms, with direct implications for implementing personalized dynamic treatments in neurology.

**HIGHLIGHTS:**

- Unsupervised learning detects enriched gene expression (GE) trajectories in disease
- These plasma and brain GE trajectories predict neuropathology and future cognitive impairment
- Most predictive molecular functions/pathways in the brain are also top predictors in the plasma
- By identifying plasma GE trajectories, patients can be easily screened and follow dynamic treatments

## INTRODUCTION

In recent decades, we have witnessed an accelerated characterization of the molecular and neuropathological mechanisms underlying neurodegenerative progression. Thanks to cutting-edge technological and methodological advances in genomic and proteomic analysis, we foresee unlimited methodological possibilities for understanding and modifying the role of genes and protein in disease (Esvelt and Wang, 2012; Mostafavi et al., 2018; Smith et al., 2016; Tan et al., 2012). Gene expression (GE) examination has been of crucial value, revealing disease-specific differentiated genes/molecular-pathways and gene-gene networks with a direct effect in neuropathological and cognitive/clinical deterioration (Mostafavi et al., 2018; Zhang et al., 2013). However, neurodegenerative conditions may take decades to develop and GE mapping techniques are quite recent, hence the unavailability of individual GE datasets covering a given disease’s whole evolution. All reported studies are based on cross-sectional or short-term longitudinal data, while we continue to lack long-term datasets covering the several phases underlying neurodegeneration.

In addition, due to its highly invasive nature, brain GE studies in neurodegeneration are based on *post-mortem* tissue samples. There are major challenges associated with the translation/extrapolation of *ex-vivo* results to *in-vivo* conditions (Ferreira et al., 2018). This could imply that disease mechanisms (e.g. gene-gene causal networks) and potential biomarkers identified with *post-mortem* data may well not be entirely generalizable to live patients. In this sense, peripheral molecular measurements (e.g. plasma GE) may be used to cross-validate *post-mortem* based methodologies and findings, potentially providing minimally invasive *in-vivo* biomarkers for accurate patient screening in the daily clinic and clinical trials implementation. Nevertheless, the lack of comprehensive longitudinal peripheral datasets, covering multiple disease stages at the individual level, makes *in-vivo* dynamic molecular analyses unpractical. Consequently, this affects the identification of robust peripheral biomarkers across continuous disease stages and variants.

Due to the proven ability to disentangle temporal components from high-dimensional cross-sectional data, novel unsupervised Machine Learning (ML) techniques offer a viable opportunity for dealing with the previous limitations. The data-driven reconstruction of pseudo-temporal paths to order observations (e.g. cells, subjects) is revolutionizing *omics* studies, enabling for the first time the mapping of complex dynamic processes using cross-sectional “snapshots” (Cannoodt et al., 2016; Gupta and Bar-Joseph, 2008; Magwene et al., 2003; Welch et al., 2016). Based on the ML inference of a low dimensional space embedded in a population’s *omics* data, and by creating a relative ordering of the individuals, we can accurately identify a series of molecular states that constitute a longitudinal trajectory for a process of interest (Campbell and Yau, 2018). When used in RNA-seq studies, this novel technique has provided an unprecedented insight into the evolution of multiple pathologies. It has also allowed tracking and dissecting differentiated spatiotemporal programs in single-cell analysis (Briggs et al., 2018).

Driven by the imperative of a better understanding and an earlier detection of neurodegeneration, here we extend pseudotemporal trajectory inference (TI) to the analysis of both *post-mortem* and *in-vivo* (blood) GE neurodegenerative samples. Firstly, to better address important methodological limitations in data exploration and visualization, we introduce the contrastive Trajectory Inference (cTI) algorithm. This allows the unsupervised identification and ordering of enriched patterns in a diseased population (e.g. Alzheimer’s and Huntington’s diseases) relative to a comparison background population (e.g. healthy elderly). Next, we analyze GE samples from blood plasma of 744 subjects in the spectrum of late-onset Alzheimer’s disease (LOAD) and from 1225 autopsied brains in the spectrum of LOAD and Huntington’s disease (HD). Our method provides molecular pathological scores that are highly predictive of neuropathological and cognitive/clinical deterioration. The results are strongly consistent for both *in-vivo* and *post-mortem* data. In addition, it allows identification of genes and molecular pathways driving neurodegenerative progression, as well as analysis of (dis)similarities in molecular disease mechanisms at brain and peripheral tissue levels. The inference of contrasted genetic trajectories is a promising tool for understanding complex neuropathological mechanisms and for minimally invasive patient screening at the daily clinic, with practical implications for implementing personalized medical interventions in neurology.

## RESULTS

### Inferring Enriched GE Neurodegenerative Trajectories

GE, neuropathology and cognitive/clinical deterioration in 1969 demented and non-demented subjects from three large-scale studies were assessed (see Figure 1, and Datasets 1-3 in *Star Methods*). GE and neuropathology evaluations from both dataset 1 (N=489, ROSMAP Study) and dataset 2 (N=736, HBTRC database) were performed in autopsied brains, with genetic profiling from the PFC. GE from Dataset 3 (N=744, ADNI database) was obtained from *in-vivo* blood samples, with all subjects also having brain imaging evaluations including amyloid PET, tau PET and/or structural MRI.

**Figure 1.**
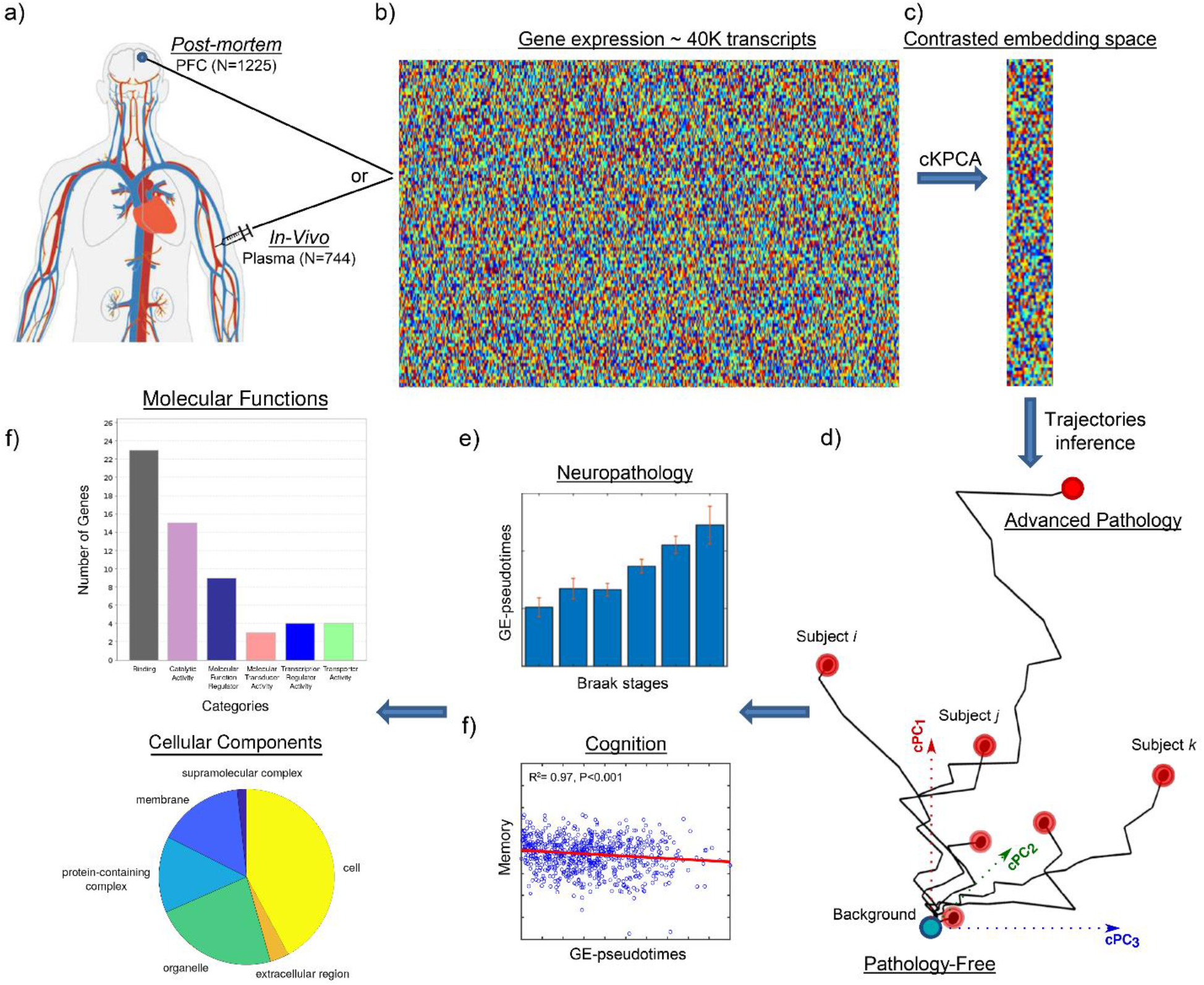
Schematic approach for GE-trajectories analysis in neurodegeneration. a) *In-vivo* blood (N=744) and *post-mortem* brain (N=1225) tissues collected. b) RNA expression for around 40,000 transcripts (dataset-specific). c) The high dimensional data is automatically reduced to an enriched space (~5 features) via a contrastive Kernel PCA algorithm (cKPCA (Abid et al., 2018)), which optimizes the exploration and visualization of the target population’s data. d) In the contrasted Principal Components (cPC) space, each subject is assigned to a GE-trajectory. The subject’s position in the corresponding GE-trajectory reflects the individual proximity to the pathology-free state (the background) and, if analyzed in the inverse direction, to the advanced disease state. An individual GE-pseudotime score is calculated, reflecting the distance to these two extremes (background or disease). e) When taken as an individual molecular score of disease evolution, the GE-pseudotime strongly predicts neuropathological and/or cognitive measurements. f) Both in peripheral and brain tissues, the cKPCA’s loadings (or weights) allow the identification and posterior functional analysis of most informative genes in terms of pathological evolution.

Aiming to uncover the molecular reconfigurations underlying neurodegenerative evolution, we proceeded to reorder the GE patterns (Fig. 1). For this, we implemented a novel unsupervised algorithm for detecting enriched trajectories in a diseased population relative to a background dataset (e.g. normal controls; see cTI subsection in *Star Methods*). A distinctive feature of cTI is the use of a contrastive Kernel PCA algorithm (Abid et al., 2018), which controls by the principal components of the background data to optimize the exploration and visualization of the target. It is a generic algorithm, adaptable to different types of data (e.g. genomic, proteomic, imaging, clinical) and nonlinear effects. Each GE dataset was firstly adjusted for relevant confounding covariates (e.g. RIN, age, gender and/or educational level; see details in *Statistics, Star Methods*). Next, the cTI was independently applied to the three populations, providing population-specific trajectories starting on the background data. Each trajectory is composed by the concatenation of a subset of subjects, which follows a given behavior in the data’s dimensionally reduced space. We hypothesized that the position of each subject in these GE trajectories would reflect individual proximity to the pathology-free state (the background) or, if analyzed in the inverse direction, proximity to advanced disease states. Correspondingly, a GE-pseudotime value ([0,1] range) is calculated for each subject, with relatively low values for subjects with final positions close to the background data, and high values for subjects on the distant extremes of the population. Notice that GE-pseudotime could then be assumed as an individual molecular score of pathological progression, whose validity is tested in the following subsections. See also Figure 1.

### Post-mortem GE Trajectories Predict Neurodegenerative Severity

Firstly, we analyzed the GE trajectories obtained for the ROSMAP study (dataset 1, N=489). The results (Figs. 2a-c) showed a clear association between the obtained molecular disease score (GE-pseudotime) and the autopsied tau and amyloid assessments, with a higher GE-pseudotime value implying an advancer neuropathologic state. Group differences in GE-pseudotime values were statistically tested via ANOVA tests with permutations. We found robust significant associations between the GE-pseudotimes and Braak stages (Fig. 2a; *F*=5.57, *P*<0.001, FEW-corrected), Cerad stages (Fig. 2b; *F*=8.39, *P*<0.001, FEW-corrected), and a composite variable (Braak+Cerad) reflecting the simultaneous presence of tau and amyloid (Fig. 2c; *F*=5.82, *P*<0.001, FEW-corrected). Notice (Fig. 2c) a clearer GE-pseudotime correspondence with the composite (Braak+Cerad) variable than with Braak and Cerad stages separately. This is in line with the concurrent accumulation of both amyloid and tau in LOAD, considered indispensable markers for characterizing this disorder’s progression (Serrano-pozo et al., 2011).

**Figure 2.**
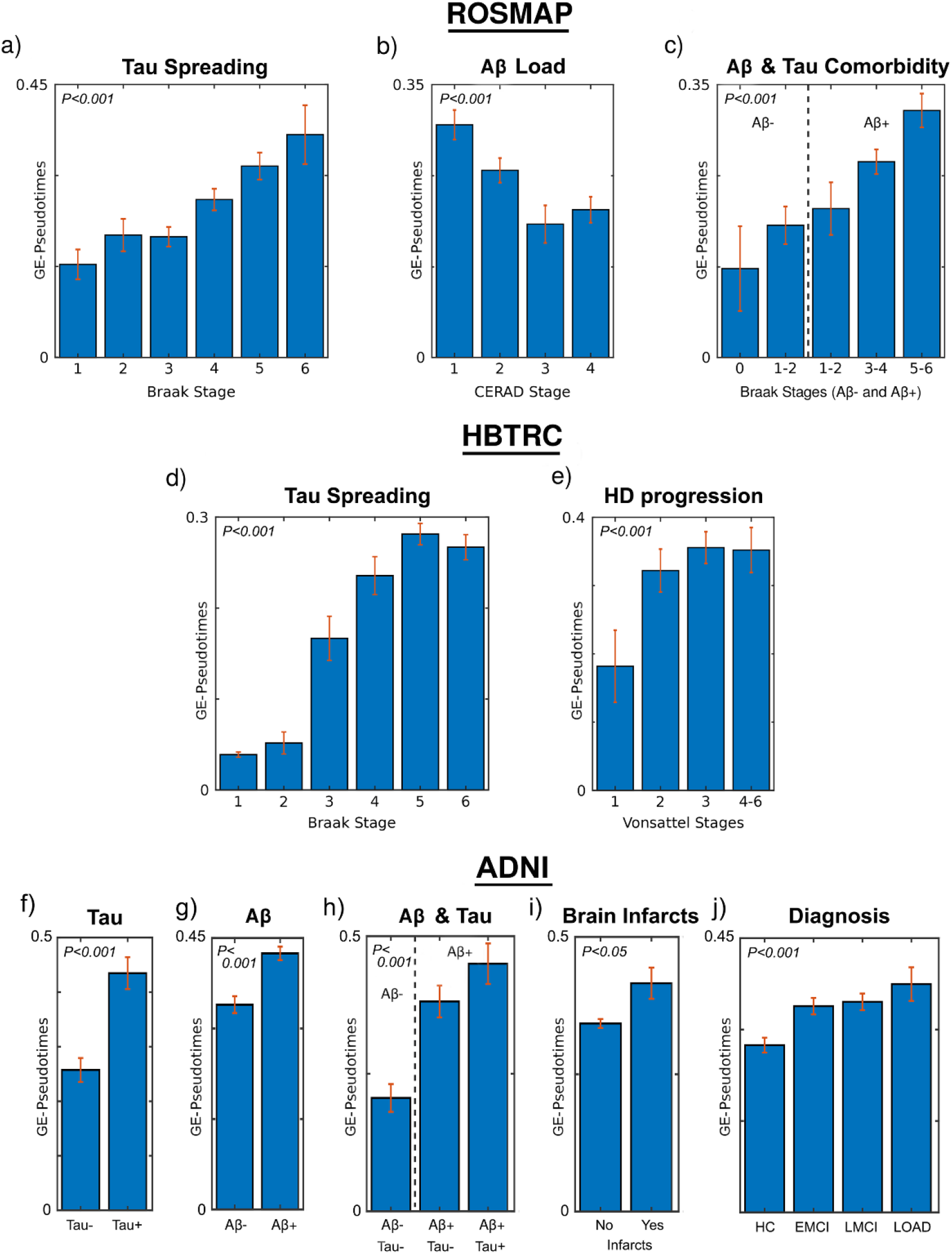
GE-based predictions of neurodegenerative severity for ROSMAP, HBTRC and ADNI populations. a-e) GE-pseudotime predictive associations with Braak (a,d), Cerad (b), Braak for Aβ-/Aβ+ (c), and Vonsattel (e) stages in ROSMAP (a-e) and HBTRC (d-e). f-i) GE-pseudotime predictive associations with Tau positivity (f), Aβ positivity, Tau-Aβ comorbidity, cerebral infarcts occurrence and clinical diagnosis in ADNI. Data is GE-pseudotime mean (± standard error). All P values are FEW-corrected (see reported values in *Results*).

Next, we explored the generalizability of these results in the considerably more heterogeneous database from HBTRC (dataset 2, N=736), including two different disorders (LOAD and HD) and nondemented controls. In consistence with the previous findings, we observed (Figs. 2d-e) a positive association between the individual molecular disease score and the levels of neuropathologic affectation in both disorders. The GE-pseudotimes were significantly associated with the Braak stages (Fig. 2d; *F*=7.87, *P*<0.001, FEW-corrected) and the Vonsattel stages (Fig. 2e; *F*= 18.80, *P*<0.001, FEW-corrected). The fact that this population included multiple disorders did not seem to affect the robustness of the subject ordering in relation with disease progression, which supports the identification of a promising biomarker for the analysis of comorbid neurological conditions.

### Blood GE as a Robust Biomarker of Neuropathological Severity and Cognitive/Clinical Deterioration

Next, we aimed to investigate if the unsupervised ordering of GE patterns present in the blood can reflect neuropathological severity and, importantly, if it could be used as a marker of present and future cognitive deterioration. If successful, the latter could have strong implications for the *in-vivo* detection of future disease evolution in the clinic and to decide if a patient should be therapeutically treated or not. To test this, we identified the enriched GE trajectories in the plasma of 744 participants in the spectrum of LOAD from ADNI (dataset 3), taking as background those subjects without cognitive/clinical alterations or any evidence of cerebral infarcts, amyloid or tau deposition (see amyloid/tau PET imaging and neuropathology evaluation for *Dataset 3*, in *Star Methods*).

In line with our previous findings with the ROSMAP and HBTRC *post-mortem* data, the ADNI-based results (Figs. 2f-j) showed a significant predictive power of pathological severity. The individual GE-pseudotime values vastly reflected the differences in tau positivity (Fig. 3f; *F*=22.12, *P*<0.001, FEW-corrected), amyloid positivity (Fig. 3g; *F*=23.03, *P*<0.001, FEW-corrected), tau-amyloid comorbidity (Fig. 3h; *F*=26.38, *P*<0.001, FEW-corrected) and brain infarcts (Fig. 3i; *F*=5.65, *P*<0.05, FEW-corrected). In addition, we tested if the identified subject ordering based on enriched GE patterns was predictive of the individual clinical and cognitive properties (Figs. 2j and 3a-d). We observed that the molecular disease score values were significantly associated with the individual clinical diagnosis (Fig. 2j; *F*=7.46, *P*<0.001, FEW-corrected). Also, they predicted 97% of the population variance in the memory (MEM) performance (Fig. 3a; R^2^_adj_=0.97, P<0.001) and, notably, a similar variance of MEM’s future rate of change in an average period of 5.35 years (Fig. 3b; R^2^_adj_=0.97, P<0.05). Interestingly, this GE-based disease progression variable showed a considerably lower predictive power for executive function (EF) in the participants, although still presenting a significant association (Fig. 3c; R^2^_adj_=0.008, P<0.05). However, it predicted around 98% of the population variance in EF’s future rate of change (Fig. 3d; R^2^_adj_=0.98, P=0.06). Altogether, these results support that, in the context of LOAD and the ADNI population, the subject’s temporal ordering based on enriched blood GE patterns is strongly reflective of neuropathological, clinical, and memory severity, being also a solid early predictor of future memory and executive function loss. It is, however, a considerably less powerful predictor of cross-sectional cognitive control.

**Figure 3.**
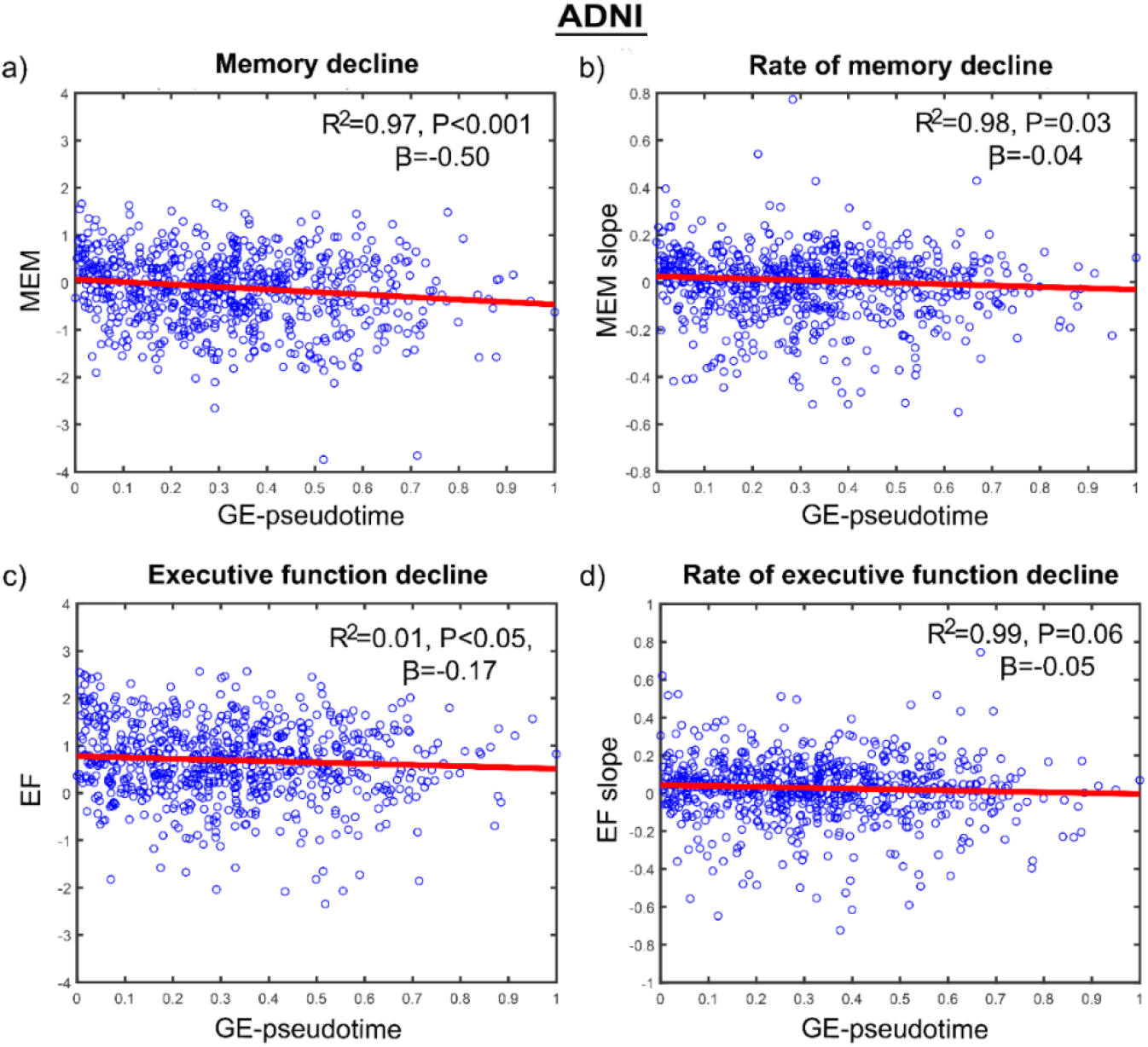
Blood GE-based predictions of Cognitive Deterioration for ADNI data. a-d) Scatter plots showing negative associations between molecular disease progression (reflected in GE-pseudotime) and measurements of cognitive integrity: MEM (a), future slope in MEM (b), EF (c) and future slope in EF (d).

### In-Vivo and Post-Mortem Molecular Pathways Underlying LOAD Progression

Next, we aimed to identify the genes, molecular functions and pathways responsible for the accurate prediction of neurodegenerative progression. We also intended to clarify if similar predictive mechanisms were common to the periphery (blood) and brain tissues. In this context, the GE-cTI can provide a quantitative mapping of the most influential genes during the process of diseased trajectories inference. Specifically, the cKPCA’s loadings (or weights) reflect how much each specific gene, in the original high dimensional space (i.e. ~40K transcripts), contributed to the reduced low dimensional space from which the trajectories were obtained. Thus, we used these weights to select the genes most influential on the subject’s ordering, i.e. those genes driving the observed population differences predictive of neuropathological and cognitive/clinical alterations across the disease’s evolution (see *Statistical Analysis*, *Star Methods*). Based on the dataset-specific identified genes, we then performed large-scale gene functional analyses with the Protein Annotation Through Evolutionary Relationship (PANTHER) classification system (Mi et al., 2013). Of note, this analysis was restricted to ROSMAP and ADNI populations, mostly related to LOAD evolution (unlike the HBTRC database, also including HD patients).

For the ROSMAP brains, we found 87 highly influential genes with 26 functional pathways (Figs. 4a,c, and Tables S2-3). These GO overrepresented pathways were highly sensitive for the detection of biological processes that are commonly associated with neuropathological and cognitive deterioration mechanisms, including oxidative stress response, axon guidance, histamine H1 receptor mediation, angiogenesis, inflammation mediated by chemokine and cytokine signaling, Wnt and VEGF signaling, apoptosis, and Alzheimer’s disease-amyloid secretase. Notably, 80% of these 26 highly predictive molecular pathways in the neurodegenerating brain (ROSMAP) were also among the most relevant pathways detected in the blood data (ADNI). The common blood-brain functional pathways relevant for LOAD progression included gastrin and cholecystokinin (CCKR) signaling, platelet derived growth factor (PDGF) signaling, B cell activation, angiogenesis, Wnt signaling, vascular endothelial growth factor (VEGF) signaling, among others (Fig. 4c and Table S4).

**Figure 4.**
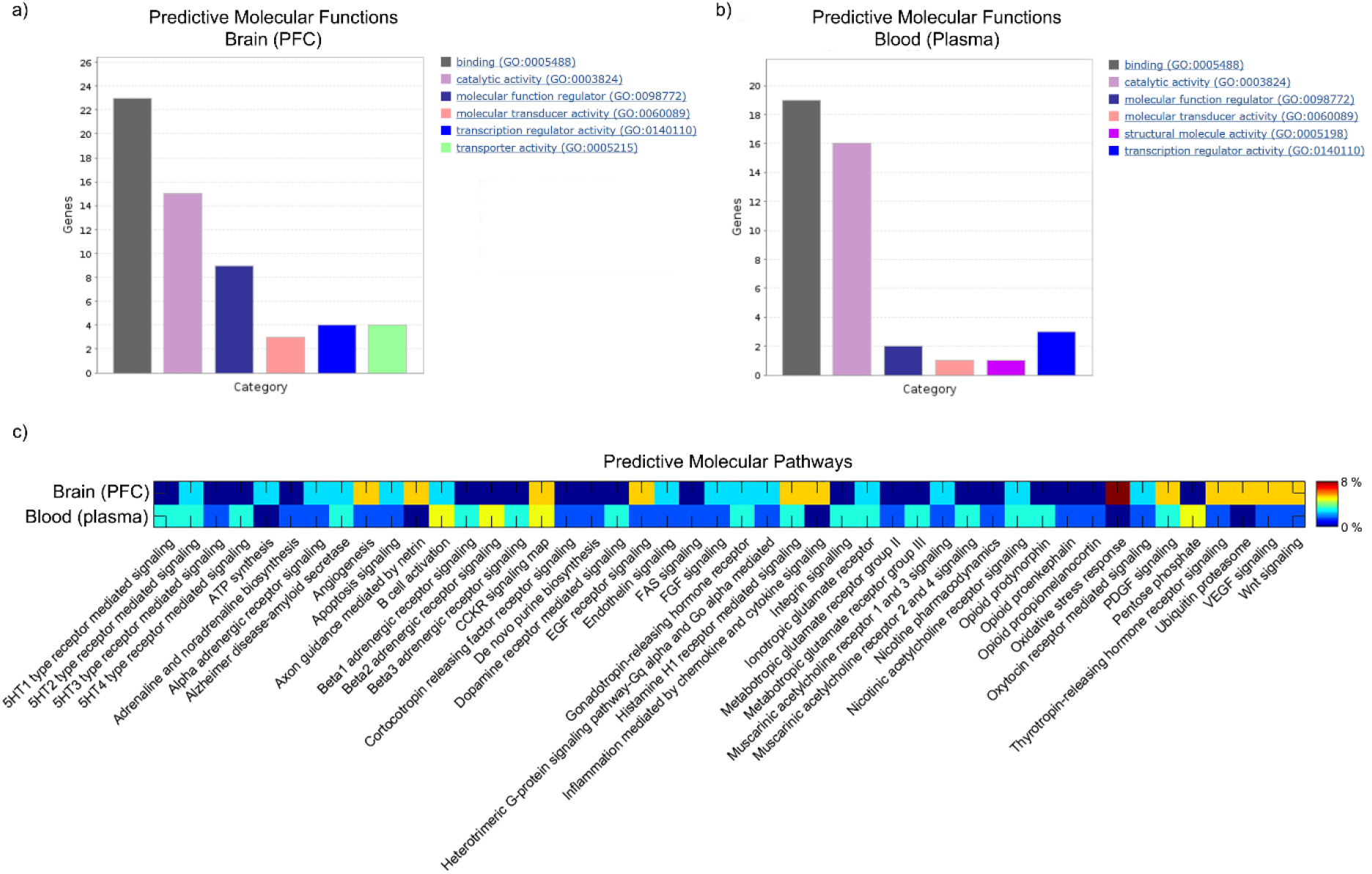
Ontology analysis of top predictor genes of LOAD development. Significant molecular functions (a,b) and pathways (c) identified in the brain’s PFC (ROSMAP) and the blood plasma (ADNI). In (a-b), the bars show the number of genes associated to the main GO overrepresented categories. In (c), the color scale indicates the presence level (in %) of each functional pathway (e.g. dark blue for absent pathways, red for highly represented pathways). For lists of genes and pathways, see Tables S2-5.

Common functional pathways CCKR (associated with appetite control and body weight regulation (Perry and Wang, 2012)), PDGF (with a significant role in blood vessel formation (Amaral et al., 2018), B cell activation (involved in immune system response), and VEGF and angiogenesis (linked to the formation of new blood vessels), evidenced the direct relationship between the central nervous system and the body, both in health and in disease. Their unsupervised data-driven identification is, therefore, supporting the crucial importance of studying the periphery-brain axis (e.g. bidirectional gut, immune and vascular interactions with brain integrity) for a better understanding of systemic pathological mechanisms underlying neurodegeneration. Interestingly, we also found another 20 highly influential molecular pathways in the blood that were not identified in the brain (Figs. 4b,c and Tables S2,5). This finding may be associated with multiple causes, including an increased pathological comorbidity in the periphery relative to the brain and/or crucial methodological limitations (i.e. the analysis of two different populations with divergent disease characteristics, and the use of different GE mapping techniques with dissimilar sensitivity/specificity capacities).

## Discussion

Due to the typically long developing period of most prevalent neurodegenerative disorders, we lack exhaustive longitudinal datasets covering the continuous molecular transitions underlying disease progression. Consequently, almost all our knowledge of the subjacent pathological mechanisms is based on data “snapshots” taken and analyzed at a few disease stages. Here, we aimed to overcome this crucial gap by inferring the intrinsic temporal information contained in large-scale neurodegenerative datasets. For that, we implemented a novel pattern analysis method that detects enriched GE trajectories in a diseased population (e.g. subjects progressing towards dementia) relative to a background population (e.g. a clinically normal control group). Our results in three different GE datasets (ROSMAP, HBTRC, ADNI) support the strong predictive power of this technique for identifying individual neuropathological stages and/or cognitive deterioration. This may well have broad implications for uncovering the dynamic mechanisms of molecular pathology, patient stratification in the clinic, and monitoring response to personalized treatments in neurodegeneration.

A minimally invasive molecular test for neurodegeneration could lead to better treatment and therapies (Ray et al., 2007). An additional aim of this study was to identify an *in-vivo* peripheral biomarker able to predict the individual’s pathophysiology and cognitive decline. When tested in 744 blood samples from ADNI, the proposed GE-cTI showed a significant association with both amyloid and tau positivity (Figs. 3a-c). Furthermore, it was able to strongly predict both the current cognitive function and its future decline (Figs. 3e-h). The fact that the proposed ML model is unsupervised (i.e. the neuropathological and cognitive variables are not used to train a predictive model), guarantees absence of possible circularity or data overfitting. Consequently, we can infer that the obtained genetic trajectories and the associated GE-pseudotime values are direct measures of molecular integrity, obtained independently of phenotypic variables, and would therefore be useful as unbiased biomarkers in clinical applications.

Our analysis of most relevant molecular pathways for predicting LOAD progression revealed a striking similarity between peripheral and intra-brain pathological mechanisms. Eighty percent of the most predictive molecular pathways identified in the brain were also identified as top predictors in the blood. These pathways support the importance of studying the peripheral-brain axis (see *Results*, last subsection), providing further evidence for a key role of gut-brain interactions (Mayer et al., 2014; Westfall et al., 2017), vascular structure and functioning (Bell and Zlokovic, 2009; Iturria-Medina et al., 2017, 2016), and immune system response (Gendelman, 2002; Labzin et al., 2018; Streit et al., 2004). The multi-tissue analysis based on genetic trajectories may be particularly useful for clarifying both local (tissue-specific) and systemic (inter-organs) neurodegenerative mechanisms.

Our method built on the pseudotemporal trajectory inference field (Cannoodt et al., 2016; Gupta and Bar-Joseph, 2008; Magwene et al., 2003; Welch et al., 2016). Modeling the dynamics of gene regulation, rather than focusing on static time points, is crucial for clarifying cellular transitions and what goes wrong in the case of disease (Cannoodt et al., 2016). We attempted to extend previous models by incorporating the use of a novel contrastive dimensionality reduction technique (Abid et al., 2018), which allows detecting enriched patterns in the population of interest while adjusting by confounding components in the background population (e.g. concurrent aging effects). In a set of complementary analyses (data not known), we observed that, in comparison with other state-of-the-art TI methods (Campbell and Yau, 2018; Welch et al., 2016), this extension provides a considerably higher sensitivity to detect diseased GE components (i.e. other methods could not predict neuropathology, nor cognition). In addition to uncovering disease dynamics, cTI may enable the data-driven identification of new subpopulations within a heterogeneous neurodegenerative population (Cannoodt et al., 2016; Trapnell, 2015; Trapnell et al., 2014), with strong implications for precision medicine and the selective enrollment of patients in clinical trials. Furthermore, once the data is ordered, it could also improve the inference of causative regulatory interactions underlying a disorder (Cannoodt et al., 2016).

Another advantage of cTI (and TI in general) is the ability to deal with high dimensional data. This is a key feature for the concurrent analysis of *multi-omics*, potentially allowing the exploration of multiple and complementary modalities, such as transcriptomics, proteomics, metabolomics and epigenomics. Contrastive trajectory inference can be also applied to the analysis of data from other fields, including multi-modal brain imaging, environmental and cognitive/clinical information. Finally, although our study is focused on neurodegenerative evolution, in general, cTI can be applicable to the study of multiple neurological and neuropsychiatric conditions.

## STAR METHODS

### Study Participants

This study used GE data (N_total_=1969) from three large-scale databases (see Table S1 for demographic characteristics). Each dataset was processed and analyzed independently:

**Dataset 1**. RNA expression data from the prefrontal cortex (PFC) of a subset of 489 autopsied subjects were downloaded from the Religious Orders Study (ROS; (Bennett et al., 2012a)) and the Memory and Aging Project Study (MAP; (Bennett et al., 2012b)). This data (Bennett et al., 2018) is available at the Accelerating Medicines Partnership Alzheimer’s Disease (AMP-AD) knowledge portal (https://www.synapse.org/#, Synapse ID 3800853). ROS (Bennett et al., 2012a) and MAP (Bennett et al., 2012b). are longitudinal clinical-pathologic cohort studies of aging, Alzheimer’s disease (AD) and related disorders. Enrollment required no known sign of dementia. Upon death, a *post-mortem* neuropathologic evaluation is performed that includes a uniform structured assessment of AD pathology, cerebral infarcts, Lewy body disease, and other pathologies common in aging and dementia. The pathologic diagnosis of AD uses NIA-Reagan and modified CERAD criteria, and the staging of neurofibrillary pathology uses Braak Staging (Braak H, 1991). An RNA integrity (RIN) score >5 and a quantity threshold (5 mg) for each sample were required (Bennett et al., 2014). cRNA was hybridized to Illumina HT-12 Expression Bead Chip (48,803 transcripts) via standard protocols using an Illumina Bead Station 500GX (Webster et al., 2009; Zhang et al., 2013).

**Dataset 2**. 736 individual *post-mortem* tissue samples from the dorsolateral prefrontal cortex BA9 of LOAD patients (N=376), HD patients (N=184) and nondemented subjects (N=173) were collected and analyzed (Zhang et al., 2013). All autopsied brains were collected by the Harvard Brain Tissue Resource Center (HBTRC; GEO accession number GSE44772), and include subjects for whom both the donor and the next of kin had completed the HBTRC informed consent (http://www.brainbank.mclean.org/). Correspondingly, tissue collection and the research were conducted according to the HBTRC guidelines (http://www.brainbank.mclean.org/). Postmortem interval (PMI) was 17.8 ± 8.3 hr, sample pH was 6.4 ± 0.3 and RNA integrity number (RIN) was 6.8 ± 0.8 for the average sample in the overall cohort.

As previously described in (Zhang et al., 2013), RNA preparation and array hybridizations applied custom microarrays manufactured by Agilent Technologies consisting of 4,720 control probes and 39,579 probes targeting transcripts representing 25,242 known and 14,337 predicted genes. Arrays were quantified on the basis of spot intensity relative to background, adjusted for experimental variation between arrays using average intensity over multiple channels, and fitted to an error model to determine significance (Emilsson et al., 2008). Braak stage, general and regional atrophy, gray and white matter atrophy and ventricular enlargement were assessed and cataloged by pathologists at McLean Hospital (Belmont, MA, USA). In addition, the severity of pathology in the HD brains was determined using the Vonsattel grading system (Vonsattel et al., 1985).

**Dataset 3**. This study used a total of 744 individual data with blood GE information, from the Alzheimer’s Disease Neuroimaging Initiative (ADNI) (adni.loni.usc.edu). The participants underwent multimodal brain imaging evaluations, including amyloid PET, tau PET and/or structural MRI. The ADNI was launched in 2003 as a public-private partnership, led by Principal Investigator Michael W. Weiner, MD. The primary goal of ADNI has been to test whether serial magnetic resonance imaging (MRI), positron emission tomography (PET), other biological markers, and clinical and neuropsychological assessments can be combined to measure the progression of mild cognitive impairment (MCI) and early Alzheimer’s disease (AD).

The Affymetrix Human Genome U219 Array (www.affymetrix.com) was used for gene expression profiling from blood samples. Peripheral blood samples were collected using PAXgene tubes for RNA analysis (Saykin et al., 2015). The quality-controlled GE data includes activity levels for 49,293 transcripts. All the participants were characterized cognitively using the mini-mental state examination (MMSE), a composite score of executive function (EF), a composite score of memory integrity (MEM) (Gibbons et al., 2012), and Alzheimer’s Disease Assessment Scale-Cognitive Subscales 11 and 13 (ADAS-11 and ADAS-13, respectively). Also, they were clinically diagnosed at baseline as healthy control (HC), early mild cognitive impairment (EMCI), late mild cognitive impairment (LMCI) or probable Alzheimer’s disease patient (LOAD).

^18^F-AV-45 (amyloid specific) and ^18^F-AV-1451 (tau specific) PET images were acquired for a subset of 660 and 166 subjects, respectively. Both amyloid and tau images were preprocessed by the Jagust Lab (UC Berkeley, US; Jagust et al., 2010). Using the amyloid images, subjects were categorized as amyloid positive (Aβ+) or negative (Aβ-) by applying a cutoff of 1.11 to a Florbetapir composite SUVR normalized by the whole cerebellum reference (Described in ADNI_UCBERKELEY_AV45_Methods_12.03.15.pdf file, ADNI database). Also, individual Freesurfer-defined cortical and subcortical brain regions were used to calculate weighted Flortaucipir averages for each region, which were normalized by the weighted Flortaucipir at the cerebellum (Described in UCBERKELEY_AV1451_Methods_Aug2018.pdf file, ADNI database). Based on the lobar classification topographic staging scheme for tau PET and the corresponding cutoff values proposed by (Schwarz et al., 2018), the subjects were staged in Braak 0 (no tau), Braak I/II, Braak III/IV or Braak V/VI. Subsequently, they were categorized as tau negative (tau-) or positive (tau+) if they were in the stages 0 or I-VI, respectively. Structural MRI images for 741 subjects were analyzed by a physician specially trained in the detection of MRI infarcts. The presence of MRI infarction was determined from the size, location and imaging characteristics of the lesion, with only lesions 3mm or larger qualifying for consideration as cerebral infarcts (Described in ADNI_UCD_MRI_Infarct_Assessment_Method_201130609.pdf file, ADNI database). Finally, a subset of subjects (N=30) was evaluated for pathological brain lesions after death. Pathological lesions were assessed using established neuropathologic diagnostic criteria (Described in ADNI_Methods_Neuropathology_Core_03-06-2018-2.pdf file, ADNI database). The analysis included histopathologic assessments of amyloid β deposits, staging of neurofibrillary tangles, scoring of neuritic plaques and assessments of co-morbid conditions such as Lewy body disease, vascular brain injury, hippocampal sclerosis, and TAR DNA binding protein (TDP) immunoreactive inclusions (Montine et al., 2012).

### Contrastive Trajectories Inference (cTI)

Given a multi-dimensional population dataset, the inference of contrasted pseudotemporal trajectories (and an individual pseudotime value) consists of four main steps:

i. For high-dimensional datasets (e.g. ~40,000 transcripts), initial selection of features most likely to be involved in a trajectory across the entire population. We apply the unsupervised method proposed by (Welch et al., 2016), which does not require prior knowledge of features involved in the process or differential expression analysis. Features are scored by comparing sample variance and neighborhood variance. A threshold is applied to select those features with higher score, e.g. we kept the features with at least a 0.95 probability of being involved in a trajectory (i.e. ~3000 gene transcripts).
ii. Data exploration and visualization via contrastive Kernel Principal Component Analysis (cKPCA; (Abid et al., 2018)). This novel technique identifies nonlinear low-dimensional patterns that are enriched in a target dataset (e.g. a diseased population) relative to a comparison background dataset (e.g. demographically matched healthy subjects). By controlling the effects of characteristic patterns in the background (e.g. pathology-free and spurious associations, noise), cKPCA (and its linear version cPCA (Abid et al., 2018)) allows visualizing specific data structures missed by standard data exploration and visualization methods (e.g. PCA, Kernel PCA). When applied to the selected GE transcripts (from step *i*), for each population, we obtained around 4-5 contrasted principal components capturing the most enriched pathological properties relative to the background (i.e. subjects without cognitive deterioration and neuropathological signs).
iii. Subjects ordering and GE-pseudotime calculation according to their proximity to the background population in the contrasted Principal Components space. For this, we first calculate the Euclidean Distance Matrix among all the subjects and the associated Minimum Spanning Tree (MST). The MST is then used to calculate the shortest trajectory/path from any subject to the background subjects. Each specific trajectory consists of the concatenation of relatively similar subjects, with a given behavior in the data’s dimensionally reduced space. The position of each subject in his/her corresponding shortest trajectory reflects the individual proximity to the pathology-free state (the background) and, if analyzed in the inverse direction, to advanced disease state. Thus, to quantify the distance to these two extremes (background or disease), an individual GE-pseudotime score is calculated as the shortest distance value to the background’s centroid, relative to the maximum population value (i.e. values are standardized between 0 and 1). Finally, the subjects are ordered according to their GE-pseudotime values, from low (close to the background group) to high values (close to the most diseased subjects).

### Statistics

Genes activity was adjusted for relevant covariates using robust linear models (Street et al., 1988). Specifically, Dataset 1 GE was adjusted for postmortem interval (PMI) in hours, age, gender and educational level. Dataset 2 GE was adjusted for PMI, sample pH, RNA integrity number (RIN), age and gender. Dataset 3 GE was controlled for RIN, Plate Number, age, gender and educational level. All predictive associations between grouping variables (e.g. Braak stages, Cerad stages, clinical diagnosis) and the individual GE-pseudotimes (see first and second *Results* subsections) were tested via ANOVA tests, FEW-controlled by permutations (Legendre and Legendre, 1998). For each dataset, a gene’s contribution to the obtained genetic trajectories was quantified as the mean of its absolute cKPCA loadings/weights. We considered 2 standard deviations over the mean for detecting most influential genes.

## Data and Code availability

The three datasets used in this study are available at the AMP-AD knowledge portal (https://www.synapse.org/#, Synapse ID 3800853), the Gene Expression Omnibus (GEO accession number GSE44772) and the ADNI database (www.adni.loni.usc.edu), respectively. We anticipate that the code will be released soon as part of an open-access software. For more information, please contact the corresponding author.

## Acknowledgments

This research was undertaken thanks in part to funding from the *Canada First Research Excellence Fund*, awarded to McGill University for the *Healthy Brains for Healthy Lives Initiative*. Dataset-1 (ROSMAP) was provided by the Rush Alzheimer’s Disease Center, Rush University Medical Center, Chicago. Dataset-1 collection was supported through funding by NIA grants P30AG10161, R01AG15819, R01AG17917, U01AG46152, and the Illinois Department of Public Health. ROSMAP data can be requested at www.radc.rush.edu. In addition, dataset-3 collection and sharing for this project was funded by ADNI (National Institutes of Health Grant U01 AG024904) and DOD ADNI (Department of Defense award number W81XWH-12-2-0012). ADNI is funded by the National Institute on Aging, the National Institute of Biomedical Imaging and Bioengineering, and through generous contributions from the following: AbbVie, Alzheimer’s Association; Alzheimer’s Drug Discovery Foundation; Araclon Biotech; BioClinica, Inc.; Biogen; Bristol-Myers Squibb Company; CereSpir, Inc.; Eisai Inc.; Elan Pharmaceuticals, Inc.; Eli Lilly and Company; EuroImmun; F. Hoffmann-La Roche Ltd and its affiliated company Genentech, Inc.; Fujirebio; GE Healthcare; IXICO Ltd.; Janssen Alzheimer Immunotherapy Research & Development, LLC.; Johnson & Johnson Pharmaceutical Research & Development LLC.; Lumosity; Lundbeck; Merck & Co., Inc.; Meso Scale Diagnostics, LLC.; NeuroRx Research; Neurotrack Technologies; Novartis Pharmaceuticals Corporation; Pfizer Inc.; Piramal Imaging; Servier; Takeda Pharmaceutical Company; and Transition Therapeutics. The Canadian Institutes of Health Research is providing funds to support ADNI clinical sites in Canada. Private sector contributions are facilitated by the Foundation for the National Institutes of Health (www.fnih.org). The grantee organization is the Northern California Institute for Research and Education, and the study is coordinated by the Alzheimer’s Disease Cooperative Study at the University of California, San Diego. ADNI data are disseminated by the Laboratory for Neuro Imaging at the University of Southern California.

## Author Contributions

ROSMAP, HBTRC and ADNI acquired the data. YIM conceived the study, implemented the programming source code, preprocessed and analyzed the data, and wrote the draft manuscript. AFK and QA prepared the figures and corrected the manuscript. All authors contributed to constructive discussions.

## SUPPLEMENTARY INFORMATION

**Table S1.** Main demographic characteristics for the three populations.

| Variable | ROSMAP<br>(N=489) | HBTRC<br>(N=736) | ADNI<br>(N=744) |
| --- | --- | --- | --- |
| Women | 299 (61.1%) | 354 (47.9%) | 336 (45.1%) |
| Age (years) | 86.1 (4.81) | 70.8 (15.10) | 73.1 (7.03) |
| Education (years) | 16.7 (3.59) | - | 16.1 (2.77) |

Data are number (%) or mean (std).

**Table S2.** Most predictive genes of LOAD progression in brain (ROSMAP) and blood (ADNI) tissues.

| Data | Genes |
| --- | --- |
| ROSMAP | METRNL, RANBP6, GPR89A, HEATR5B, DUSP12, EPYC, CTSL1, HS.66187, BAZ1A, WWTR1, GIMAP8, ZFAND3, PRKCE, ASB6, LETMD1, C3ORF59, ZNF667, SPOCD1, ABLIM2, RARRES1, LANCL1, PLCG2, TNNC1, TATDN1, DCLK1, TNRC6B, BPGM, AMACR, TAC1, MYO9B, UBE2W, KCNIP4, C3ORF38, NR1H3, TSPYL1, C6ORF170, WDR61, ATP1B1, PSMC2, CTSL1, LOC220686, EVI5, HSPB7, BRMS1, LARGE, SLC7A7, CAMK2A, KIAA1688, RAB11FIP2, YME1L1, RAP1GAP, MAP1A, SETBP1, PSMD10, LMO3, SNTG1, ZDHHC3, MMD TRIM32, CMTM3, FAM40B, ATXN10, CYCS, DNAJC12, SIL1, HSD17B11, NAP1L5, ATP6V1E1, TAC1, DCLRE1C, PAP2D, MFSD4, DNHL1, ZNF223, PGM2L1, RBM11, HECTD1, TSC22D1, CDKL2, NUDT11, MOBKL2C, TGFBI |
| ADNI | G6PD, C11ORF68, SETD5, ENOPH1, SOD1, EDC4, ZSWIM8, ACADSB, ACAP2, VPS72, SFSWAP, ZNF302, ZBED5, OS9, SMARCC2, DMTF1, DAXX, STX3, BRD2, RTN3, SERP1, SERP1, HNRNPUL2, SMNDC1, UBN1, SLC6A6, PCNX, MIDN, CAPZA2, PGD, CHD7, AGPS, DYRK2, WDR44, WDR44, ABI1, PDLIM7, EIF4A2, CFP, PRKCD, NFAM1, ABI1, SDCBP, NADK, PNPLA8, ARHGAP26, SDCBP, SBDSP1, SBDS, ABI1, KDM5A, S1PR1, CHD8, BRMS1, PRRC2C, SSFA2, ANXA11, RASSF2, ANXA11, VTA1, PAF1, TNIP1, GBA, GIGYF2, OS9, CHST15, GARS, SDCBP, ARGLU1, PPP1CB, COPS6, VAMP7, SNAP23, CNOT6, SDHD, SDCBP, CREBRF, PDLIM7, DOCK5, SMNDC1 |

**Table S3.** Molecular pathways underlying GE trajectories associated to LOAD progression in brain tissues (ROSMAP).

| Pathway name | Presence (%) |
| --- | --- |
| Oxidative stress response | 7.7 |
| Axon guidance mediated by netrin | 5.1 |
| Histamine H1 receptor mediated signaling pathway | 5.1 |
| Angiogenesis | 5.1 |
| Inflammation mediated by chemokine and cytokine signaling pathway | 5.1 |
| Ubiquitin proteasome pathway | 5.1 |
| EGF receptor signaling pathway | 5.1 |
| PDGF signaling pathway | 5.1 |
| Thyrotropin-releasing hormone receptor signaling pathway | 5.1 |
| CCKR signaling map | 5.1 |
| Wnt signaling pathway | 5.1 |
| VEGF signaling pathway | 5.1 |
| ATP synthesis | 2.6 |
| Apoptosis signaling pathway | 2.6 |
| Ionotropic glutamate receptor pathway | 2.6 |
| Alzheimer disease-amyloid secretase pathway | 2.6 |
| Alpha adrenergic receptor signaling pathway | 2.6 |
| Endothelin signaling pathway | 2.6 |
| Gonadotropin-releasing hormone receptor pathway | 2.6 |
| Nicotinic acetylcholine receptor signaling pathway | 2.6 |
| Oxytocin receptor mediated signaling pathway | 2.6 |
| Muscarinic acetylcholine receptor 1 and 3 signaling pathway | 2.6 |
| B cell activation | 2.6 |
| Heterotrimeric G-protein signaling pathway-Gq alpha and Go alpha mediated pathway | 2.6 |
| 5HT2 type receptor mediated signaling pathway | 2.6 |
| FGF signaling pathway | 2.6 |

**Table S4.** Common molecular pathways underlying GE trajectories associated to LOAD progression in blood (ADNI) and brain (ROSMAP) tissues.

| **Pathway name** | **Presence (%)** |
| --- | --- |
| CCKR signaling map | 4.85 |
| PDGF signaling pathway | 4.10 |
| Histamine H1 receptor mediated signaling pathway | 4.10 |
| B cell activation | 3.60 |
| Angiogenesis | 3.30 |
| Wnt signaling pathway | 3.30 |
| VEGF signaling pathway | 3.30 |
| Thyrotropin-releasing hormone receptor signaling pathway | 3.30 |
| EGF receptor signaling pathway | 3.30 |
| Ionotropic glutamate receptor pathway | 2.85 |
| 5HT2 type receptor mediated signaling pathway | 2.85 |
| Alzheimer disease-amyloid secretase pathway | 2.85 |
| Nicotinic acetylcholine receptor signaling pathway | 2.85 |
| Gonadotropin-releasing hormone receptor pathway | 2.85 |
| Apoptosis signaling pathway | 2.05 |
| Alpha adrenergic receptor signaling pathway | 2.05 |
| Heterotrimeric G-protein signaling pathway-Gq alpha and Go alpha mediated pathway | 2.05 |
| FGF signaling pathway | 2.05 |
| Oxytocin receptor mediated signaling pathway | 2.05 |
| Endothelin signaling pathway | 2.05 |
| Muscarinic acetylcholine receptor 1 and 3 signaling pathway | 2.05 |

**Table S5.** Molecular pathways underlying GE trajectories associated to LOAD progression in blood tissues (ADNI).

| Pathway name | Presence (%) |
| --- | --- |
| Beta2 adrenergic receptor signaling pathway | 4.6 |
| Pentose phosphate pathway | 4.6 |
| B cell activation | 4.6 |
| CCKR signaling map | 4.6 |
| Beta3 adrenergic receptor signaling pathway | 3.1 |
| Metabotropic glutamate receptor group III pathway | 3.1 |
| Beta1 adrenergic receptor signaling pathway | 3.1 |
| 5HT4 type receptor mediated signaling pathway | 3.1 |
| Ionotropic glutamate receptor pathway | 3.1 |
| 5HT2 type receptor mediated signaling pathway | 3.1 |
| Alzheimer disease-amyloid secretase pathway | 3.1 |
| 5HT1 type receptor mediated signaling pathway | 3.1 |
| Integrin signalling pathway | 3.1 |
| PDGF signaling pathway | 3.1 |
| Opioid prodynorphin pathway | 3.1 |
| Histamine H1 receptor mediated signaling pathway | 3.1 |
| Nicotinic acetylcholine receptor signaling pathway | 3.1 |
| Muscarinic acetylcholine receptor 2 and 4 signaling pathway | 3.1 |
| Dopamine receptor mediated signaling pathway | 3.1 |
| Gonadotropin-releasing hormone receptor pathway | 3.1 |
| Apoptosis signaling pathway | 1.5 |
| De novo purine biosynthesis | 1.5 |
| Angiogenesis | 1.5 |
| 5HT3 type receptor mediated signaling pathway | 1.5 |
| Alpha adrenergic receptor signaling pathway | 1.5 |
| Adrenaline and noradrenaline biosynthesis | 1.5 |
| Nicotine pharmacodynamics pathway | 1.5 |
| Heterotrimeric G-protein signaling pathway-Gq alpha and Go alpha mediated pathway' | 1.5 |
| Wnt signaling pathway | 1.5 |
| VEGF signaling pathway | 1.5 |
| Thyrotropin-releasing hormone receptor signaling pathway | 1.5 |
| FGF signaling pathway | 1.5 |
| Oxytocin receptor mediated signaling pathway | 1.5 |
| FAS signaling pathway | 1.5 |
| Endothelin signaling pathway | 1.5 |
| EGF receptor signaling pathway | 1.5 |
| Opioid proopiomelanocortin pathway | 1.5 |
| Opioid proenkephalin pathway | 1.5 |
| Muscarinic acetylcholine receptor 1 and 3 signaling pathway | 1.5 |
| Corticotropin releasing factor receptor signaling pathway | 1.5 |
| Metabotropic glutamate receptor group II pathway | 1.5 |

## REFERENCES

Abid, A., Zhang, M.J., Bagaria, V.K., Zou, J., 2018. Exploring patterns enriched in a dataset with contrastive principal component analysis. Nat. Commun. 9, 1–24. doi:10.1038/s41467-018-04608-8

Amaral, R., Cavanagh, B., O’Brien, F., Kearney, C., 2018. Platelet-derived growth factor stabilises vascularisation in collagen-glycosaminoglycan scaffolds in vitro. J. Tissue Eng. Regen. Med.

Bell, R.D., Zlokovic, B. V, 2009. Neurovascular mechanisms and blood-brain barrier disorder in Alzheimer’s disease. Acta Neuropathol. 118, 103–13. doi:10.1007/s00401-009-0522-3

Bennett, D., Schneider, J., Arvanitakis, Z., Wilson, R., 2012a. Overview and findings from the religious orders study. Curr Alzheimer Res. 9, 628–645.

Bennett, D., Schneider, J., Buchman, A., Barnes, L., Boyle, P., Wilson, R., 2012b. Overview and Findings from the Rush Memory and Aging Project 9, 646–663.

Bennett, D.A., Buchman, A.S., Boyle, P.A., Barnes, L.L., Wilson, R.S., Schneider, J.A., 2018. Religious Orders Study and Rush Memory and Aging Project. J. Alzheimer’s Dis. 64, S161–S189. doi:10.3233/JAD-179939

Bennett, D.A., Yu, L., De Jager, P.L., 2014. Building a pipeline to discover and validate novel therapeutic targets and lead compounds for Alzheimer’s disease. Biochem. Pharmacol. 88, 617–630. doi:10.1016/j.bcp.2014.01.037

Braak H, B.E., 1991. Neuropathological stageing of Alzheimer-related changes. Acta Neuropathol. 82, 239–59.

Briggs, J.A., Weinreb, C., Wagner, D.E., Megason, S., Peshkin, L., Kirschner, M.W., Klein, A.M., 2018. The dynamics of gene expression in vertebrate embryogenesis at single-cell resolution. Science (80-.). 360. doi:10.1126/science.aar5780

Campbell, K.R., Yau, C., 2018. Uncovering pseudotemporal trajectories with covariates from single cell and bulk expression data. Nat. Commun. 9. doi:10.1038/s41467-018-04696-6

Cannoodt, R., Saelens, W., Saeys, Y., 2016. Computational methods for trajectory inference from single-cell transcriptomics. Eur. J. Immunol. 46, 2496–2506. doi:10.1002/eji.201646347

Emilsson, V., Thorleifsson, G., Zhang, B., Leonardson, A.S., Zink, F., Zhu, J., Carlson, S., Helgason, A., Walters, G.B., Gunnarsdottir, S., Mouy, M., Steinthorsdottir, V., Eiriksdottir, G.H., Bjornsdottir, G., Reynisdottir, I., Gudbjartsson, D., Helgadottir, A., Jonasdottir, A., Jonasdottir, A., Styrkarsdottir, U., Gretarsdottir, S., Magnusson, K.P., Stefansson, H., Fossdal, R., Kristjansson, K., Gislason, H.G., Stefansson, T., Leifsson, B.G., Thorsteinsdottir, U., Lamb, J.R., Gulcher, J.R., Reitman, M.L., Kong, A., Schadt, E.E., Stefansson, K., 2008. Genetics of gene expression and its effect on disease. Nature 452, 423–428. doi:10.1038/nature06758

Esvelt, K., Wang, H., 2012. Genome-scale engineering for systems and synthetic biology. Mol. Syst. Biol. 9, 641.

Ferreira, P.G., Muñoz-Aguirre, M., Reverter, F., Sá Godinho, C.P., Sousa, A., Amadoz, A., Sodaei, R., Hidalgo, M.R., Pervouchine, D., Carbonell-Caballero, J., Nurtdinov, R., Breschi, A., Amador, R., Oliveira, P., Çubuk, C., Curado, J., Aguet, F., Oliveira, C., Dopazo, J., Sammeth, M., Ardlie, K.G., Guigó, R., 2018. The effects of death and post-mortem cold ischemia on human tissue transcriptomes. Nat. Commun. 9. doi:10.1038/s41467-017- 02772-x

Gendelman, H.E., 2002. Neural immunity: Friend or foe? J. Neurovirol. 8, 474–9. doi:10.1080/13550280290168631

Gibbons, L.E., Carle, A.C., Mackin, R.S., Harvey, D., Mukherjee, S., Insel, P., Curtis, S.M.K., Mungas, D., Crane, P.K., 2012. A composite score for executive functioning, validated in Alzheimer’s Disease Neuroimaging Initiative (ADNI) participants with baseline mild cognitive impairment. Brain Imaging Behav. 6, 517–527. doi:10.1007/s11682-012-9176-1

Gupta, A., Bar-Joseph, Z., 2008. Extracting dynamics from static cancer expression data. IEEE/ACM Trans. Comput. Biol. Bioinforma. 5, 172–182. doi:10.1109/TCBB.2007.70233

Iturria-Medina, Y., Carbonell, F.M., Sotero, R.C., Chouinard-Decorte, F., Evans, A.C., 2017. Multifactorial causal model of brain (dis)organization and therapeutic intervention: Application to Alzheimer’s disease. Neuroimage 152, 60–77. doi:10.1016/j.neuroimage.2017.02.058

Iturria-Medina, Y., Sotero, R.C., Toussaint, P.J., Mateos-Perez, J.M., Evans, A.C., Initiative, T.A.D.N., 2016. Early role of vascular dysregulation on late-onset Alzheimer’s disease based on multifactorial data-driven analysis. Nat Commun 7, 11934. doi:10.1038/ncomms11934

Labzin, L., Heneka, M., Latz, E., 2018. Innate Immunity and Neurodegeneration. Annu. Rev. Med. 69, 437–449.

Legendre, P., Legendre, L., 1998. Numerical ecology, 2nd Englis. ed. Elsevier Science BV, Amsterdam.

Magwene, P.M., Kim, P.L., Junhyong, 2003. Reconstructing the temporal ordering of biological samples using microarray data. Bioinformatics Vol. 19, 842–850.

Mayer, E.A., Knight, R., Mazmanian, S.K., Cryan, J.F., Tillisch, K., 2014. Gut Microbes and the Brain: Paradigm Shift in Neuroscience. J. Neurosci. 34, 15490–15496. doi:10.1523/JNEUROSCI.3299-14.2014

Mi, H., Muruganujan, A., Casagrande, J.T., Thomas, P.D., 2013. Large-scale gene function analysis with the PANTHER classification system. Nat. Protoc. 8, 1551–1566. doi:10.1038/nprot.2013.092

Montine, T.J., Phelps, C.H., Beach, T.G., Bigio, E.H., Cairns, N.J., Dickson, D.W., Duyckaerts, C., Frosch, M.P., Masliah, E., Mirra, S.S., Nelson, P.T., Schneider, J.A., Thal, D.R., Trojanowski, J.Q., Vinters, H. V., Hyman, B.T., 2012. National institute on aging-Alzheimer’s association guidelines for the neuropathologic assessment of Alzheimer’s disease: A practical approach. Acta Neuropathol. 123, 1–11. doi:10.1007/s00401-011-0910-3

Mostafavi, S., Gaiteri, C., Sullivan, S.E., White, C.C., Tasaki, S., Xu, J., Taga, M., Klein, H., Patrick, E., Komashko, V., Mccabe, C., Smith, R., Bradshaw, E.M., Root, D.E., Regev, A., Yu, L., Chibnik, L.B., Schneider, J.A., Young-pearse, T.L., Bennett, D.A., Jager, P.L. De, 2018. decline of Alzheimer’ s disease. Nat. Neurosci. 21. doi:10.1038/s41593-018-0154-9

Perry, B., Wang, Y., 2012. Appetite regulation and weight control: The role of gut hormones. Nutr. Diabetes 2, e26–7. doi:10.1038/nutd.2011.21

Ray, S., Britschgi, M., Herbert, C., Takeda-Uchimura, Y., Boxer, A., Blennow, K., Friedman, L.F., Galasko, D.R., Jutel, M., Karydas, A., Kaye, J. a, Leszek, J., Miller, B.L., Minthon, L., Quinn, J.F., Rabinovici, G.D., Robinson, W.H., Sabbagh, M.N., So, Y.T., Sparks, D.L., Tabaton, M., Tinklenberg, J., Yesavage, J. a, Tibshirani, R., Wyss-Coray, T., 2007. Classification and prediction of clinical Alzheimer’s diagnosis based on plasma signaling proteins. Nat. Med. 13, 1359–62. doi:10.1038/nm1653

Saykin, A.J., Shen, L., Yao, X., Kim, S., Nho, K., Risacher, S.L., Ramanan, V.K., Foroud, T.M., Faber, K.M., Sarwar, N., Munsie, L.M., Hu, X., Soares, H.D., Potkin, S.G., Thompson, P.M., Kauwe, J.S.K., Kaddurah-Daouk, R., Green, R.C., Toga, A.W., Weiner, M.W., 2015. Genetic studies of quantitative MCI and AD phenotypes in ADNI: Progress, opportunities, and plans. Alzheimer’s Dement. 11, 792–814. doi:10.1016/j.jalz.2015.05.009

Schwarz, A.J., Shcherbinin, S., Slieker, L.J., Risacher, S.L., Charil, A., Irizarry, M.C., Fleisher, A.S., Southekal, S., Joshi, A.D., Devous, M.D., Miller, B.B., Saykin, A.J., 2018. Topographic staging of tau positron emission tomography images. Alzheimer’s Dement. s9–II, 47. doi:10.1016/j.dadm.2018.01.006

Serrano-pozo, A., Frosch, M.P., Masliah, E., Hyman, B.T., 2011. Neuropathological Alterations in Alzheimer Disease 1–23. doi:10.1101/cshperspect.a006189

Smith, A.R., Mill, J., Smith, R.G., Lunnon, K., 2016. Neuroepigenetics Elucidating novel dysfunctional pathways in Alzheimer’ s disease by integrating loci identified in genetic and epigenetic studies. NEPIG 6, 32–50. doi:10.1016/j.nepig.2016.05.001

Street, J.O., Carroll, R.J., Ruppert, D., 1988. A Note on Computing Robust Regression Estimates via Iteratively Reweighted Least Squares. Am. Stat. 42, 152–154.

Streit, W.J., Mrak, R.E., Griffin, W.S.T., 2004. Microglia and neuroinflammation: a pathological perspective. J. Neuroinflammation 1, 14. doi:10.1186/1742-2094-1-14

Tan, W., Carlson, D., Walton, M.W., Fahrenkrug, S., Hackett, P., 2012. Precision Editing of Large Animal Genomes. Adv. Genet. 80, 37–97. doi:10.1016/B978-0-12-404742-6.00002-8

Trapnell, C., 2015. Defining cell types and states with single-cell genomics. Genome Res. 25, 1491–1498. doi:10.1101/gr.190595.115

Trapnell, C., Cacchiarelli, D., Grimsby, J., Pokharel, P., Li, S., Morse, M., Lennon, N.J., Livak, K.J., Mikkelsen, T.S., Rinn, J.L., 2014. The dynamics and regulators of cell fate decisions are revealed by pseudotemporal ordering of single cells. Nat. Biotechnol. 32, 381–386. doi:10.1038/nbt.2859

Vonsattel, J., Myers, R., Stevens, T., Ferrante, R., Bird, E., Richardson, J.E.P., 1985. Neuropathological Classification of Huntington’s disease. J. Neuropathol. Exp. Neurol. 44, 559–577. doi:10.1111/j.1600-0501.2010.02096.x

Webster, J.A., Gibbs, J.R., Clarke, J., Ray, M., Zhang, W., Holmans, P., Rohrer, K., Zhao, A., Marlowe, L., Kaleem, M., McCorquodale, D.S., Cuello, C., Leung, D., Bryden, L., Nath, P., Zismann, V.L., Joshipura, K., Huentelman, M.J., Hu-Lince, D., Coon, K.D., Craig, D.W., Pearson, J. V., Heward, C.B., Reiman, E.M., Stephan, D., Hardy, J., Myers, A.J., 2009. Genetic Control of Human Brain Transcript Expression in Alzheimer Disease. Am. J. Hum. Genet. 84, 445–458. doi:10.1016/j.ajhg.2009.03.011

Welch, J.D., Hartemink, A.J., Prins, J.F., 2016. SLICER: Inferring branched, nonlinear cellular trajectories from single cell RNA-seq data. Genome Biol. 17, 1–15. doi:10.1186/s13059-016-0975-3

Westfall, S., Lomis, N., Kahouli, I., Dia, S.Y., Singh, S.P., Prakash, S., 2017. Microbiome, probiotics and neurodegenerative diseases: deciphering the gut brain axis. Cell. Mol. Life Sci. 74, 3769–3787. doi:10.1007/s00018-017-2550-9

Zhang, B., Gaiteri, C., Bodea, L.-G., Wang, Z., McElwee, J., Podtelezhnikov, A., Zhang, C., 2013. Integrated Systems Approach Identifies Genetic Nodes and Networks in LOAD 153, 707–720.

